# A long-read draft assembly of the Chinese mantis (Mantodea: Mantidae: *Tenodera sinensis*) genome reveals patterns of ion channel gain and loss across Arthropoda

**DOI:** 10.1101/2023.11.15.567226

**Authors:** JK Goldberg, RK Godfrey, M Barrett

## Abstract

Praying mantids (Mantodea: Mantidae) are iconic insects that have captivated biologists for decades, especially the species with cannibalistic copulatory behavior. This behavior has been cited as evidence that insects lack nociceptive capacities and cannot feel pain; however, this behaviorally-driven hypothesis has never been rigorously tested at the genetic or functional level. To enable future studies of nociceptive capabilities in mantids, we sequenced and assembled a draft genome of the Chinese praying mantis (*Tenodera sinensis*) and identified multiple classes of nociceptive ion channels by comparison to orthologous gene families in Arthropoda. Our assembly - produced using PacBio HiFi reads - is not chromosome-scale (Total size = 3.03Gb; N50 = 1.8Mb; 4966 contigs), but is highly complete with respect to gene content (BUSCO complete = 98.7% [odb10_insecta]). The size of our assembly is substantially larger than that of most other insects, but is consistent with the size of other mantid genomes. We found that most families of nociceptive ion channels are present in the *T. sinensis* genome; that they are most closely related to those found in the damp-wood termite (*Zootermopsis nevadensis*); and that some families have expanded in *T. sinensis* while others have contracted relative to nearby lineages. Our findings suggest that mantids are likely to possess nociceptive capabilities and provide a foundation for future experimentation regarding ion channel functions and their consequences for insect behavior.

## Introduction

The Chinese mantis, *Tenodera sinensis* (Saussure, 1871; Mantidae), is a large-bodied mantid species native to Asia and the nearby islands. In the late 1800s, the species was accidentally introduced to the Philadelphia area (Blatchley, 1920), and has become established throughout the contiguous United States (GBIF.org, 2023; Figure 1).

**Figure 1.**
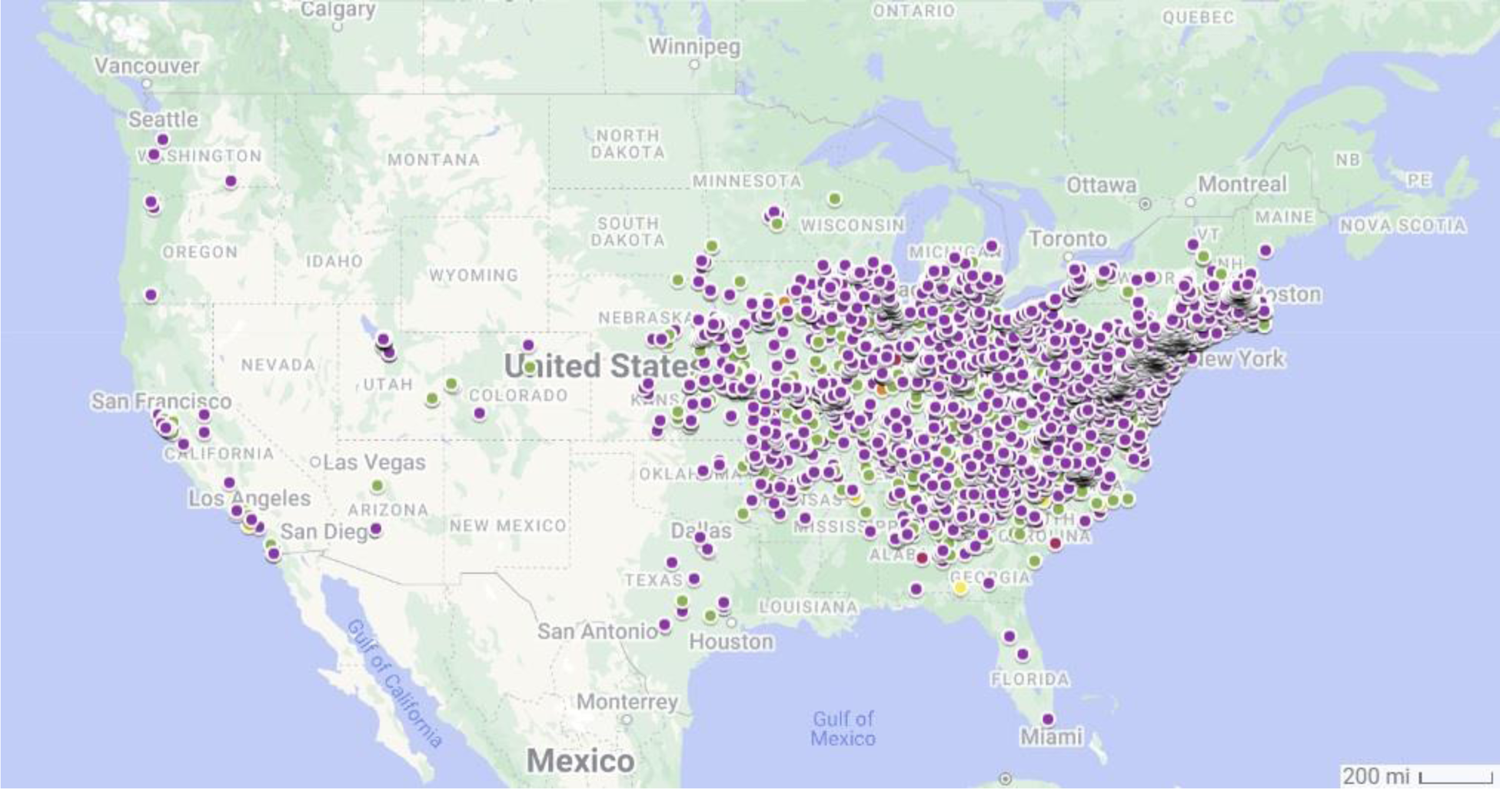
GBIF occurrence records of *T. sinensis* in the United States as of 2023. Color indicates year of record (red = pre-2000; orange = 2000-2009; yellow = 2010-2014; green = 2015-2019; purple = 2020-2023).

As a successful, invasive, generalist predator (Crowder and Snyder, 2010), *T. sinensis* has become an important ecological study species in its non-native range (Moran and Hurd 1994, Moran et al. 1996, Fagan et al. 2002). *T. sinensis* are ambush predators, spending 93% of their time waiting for prey (Rathet and Hurd, 1983). Depending on life stage and body size (Hurd et al. 2015), these mantises eat insects (including other mantises), spiders, slugs, and even hummingbirds (Hurd 1988, Iwasaki 1998, Verchot 2014, Mebs et al. 2017, Wilson Rankin et al. 2023). *T. sinensis* is probably most well-known as a study system for sexual cannibalism, where females may eat males during copulation (though field observations suggest this actually occurs in less than 10% of cases; Hurd et al. 1994), or before/after copulation. Food limitation during oogenesis is expected to drive female mantises to continuously attract mates which, if consumed, increase fecundity (O’Hara and Brown 2021; Brown and Barry 2016). Males may make up 63% of female diets during this crucial reproductive period (Hurd et al. 1994). Males are known to adjust their approach behaviors in response to the perceived threat level posed by a female and female encounter rates (Lelito and Brown, 2006, Brown et al. 2012); unlike some other sexually cannibalistic species, male *T. sinensis* are not self-sacrificial (Buskirk et al. 1984).

Despite males’ well-studied risk-avoidance behaviors that suggest a lack of complicity (see Lelito and Brown, 2006, and discussion in Christensen and Brown, 2018), sexual cannibalism in mantids has featured prominently in discussions of the plausibility of insect pain and, thus, sentience (e.g., as in Eisemann et al. 1984; though see Gibbons and Sarlak 2020). Male mantises may be unable to perceive (minimally) mechanically noxious stimuli, which animals typically perceive using nociceptive ion channels in their peripheral sensory neurons. Eisemann et al. (1984) suggest that the lack of nociceptive ability, as demonstrated by males continuing to mate while being cannibalized (e.g., behavioral non-responsiveness), precludes any plausible further possibility of pain experience. This would make mantids an interesting species for studies of insect pain perception, a field of growing research interest given the development of large-scale insect farming and associated welfare concerns (e.g., van Huis, 2021, Gibbons et al., 2022, Barrett & Adcock, 2023).

Here, we present an assembly of the Chinese praying mantis genome (*Tenodera sinensis*) and combine it with existing genomic resources to study the evolution of nociceptive channel evolution across the arthropod phylum. We find that mantids have genes that encode many well-studied arthropod nociceptors dedicated to perceiving diverse noxious stimuli (mechanical, chemical, thermal). Across the arthropods, we find that some channel families are conserved, whereas copy numbers vary greatly in others. We discuss the potential factors that may underlie this variation and the implications thereof for rearing and domestication of arthropods.

## Materials & Methods

### Mantis rearing

The specimen used for whole-genome sequencing (adult female; Figure 2) was reared at 50% humidity, 27 ℃, 14:10 L:D from a set of six ootheca purchased from Carolina Biological in spring 2023. Mantises were reared collectively until the third instar, then reared in separate containers and fed flightless *Drosophila melanogaster* and *D. hydei, Acheta domesticus* cricket nymphs, and *Tenebrio molitor* mealworm larvae. The individual used for analyses (Figure 2) was flash frozen on liquid nitrogen and sent to the Arizona Genomics Institute (AGI, Tucson, AZ, USA) for DNA extraction and sequencing.

**Figure 2.**
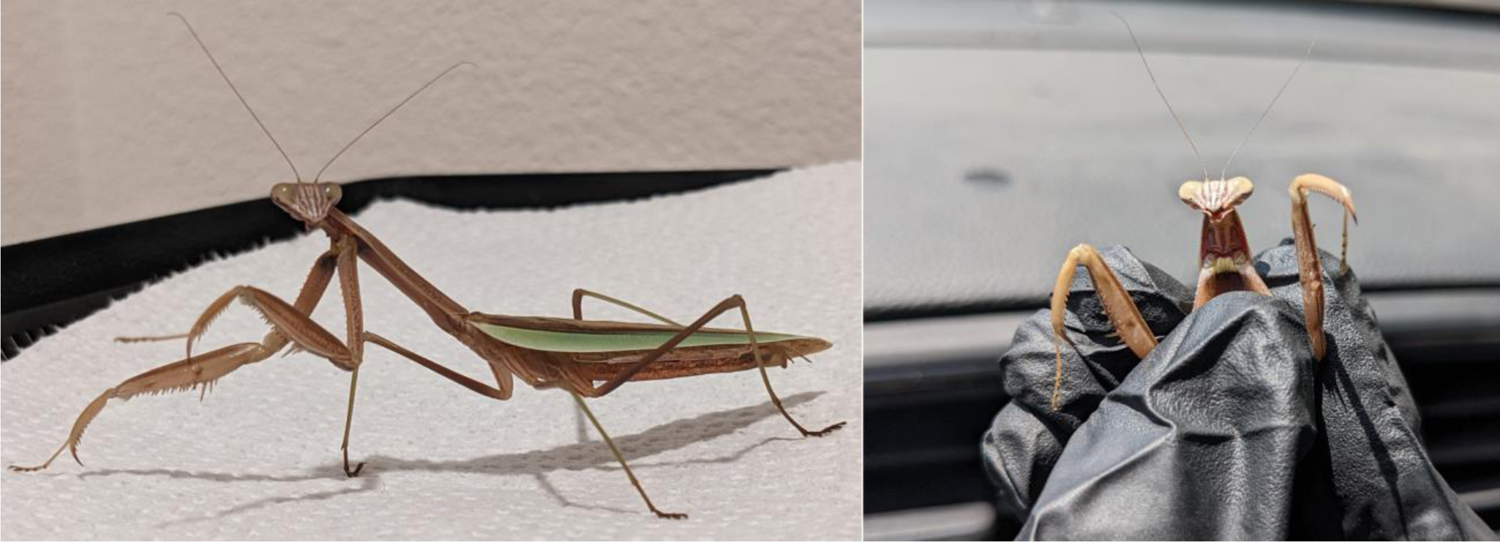
Photos of adult female *T. sinensis* specimen that was used for genome assembly.

### DNA extraction

High molecular weight DNA was extracted from ground tissue in an extraction buffer with Tris HCl buffer 0.1M pH 8.0, EDTA 0.1M pH8, SDS 1% and Proteinase K in 50C for 60 minutes. Mixture was spun down and aqueous phase transferred to a new tube. 5M Potassium acetate was added, precipitated on ice, and spun down. After centrifugation, the supernatant was gently extracted with 24:1 chloroform:isoamyl alcohol. The upper phase was transferred to a new tube and DNA precipitated with iso-propanol. DNA was collected by centrifugation, washed with 70% ethanol, air dried and dissolved thoroughly in 1x TE followed by RNAse treatment. DNA purity was measured with Nanodrop, DNA concentration measured with Qubit HS kit (Invitrogen) and DNA size was validated by Femto Pulse System (Agilent).

### Genome sequencing

DNA was sheared to an appropriate size range (10-20 kb) using Megaruptor 3 (Diagenode) followed by SMRTbell cleanup beads. The sequencing library was constructed following manufacturers protocols using SMRTbell Prep kit 3.0. The final library was size selected on a Pippin HT (Sage Science) using S1 marker with a 10-25 kb size selection. The recovered final library was quantified with Qubit HS kit (Invitrogen) and size checked on Femto Pulse System (Agilent). The final library was prepared for sequencing with PacBio Sequel II Sequencing kit 3.1 for HiFi library, loaded on Revio SMRT cells, and sequenced in CCS mode for 24 hours.

### Genome assembly and annotation

CCS output (ie: HiFi reads; 6174839 reads; 95 Gb Q32; mean length = 15396) were assembled using hifiasm-0.16.0 (Cheng et al. 2021; RRID:SCR_021069). Assembly was visualized using Bandage v0.8.1 (Wick et al. 2015; RRID:SCR_022772), which also provided contiguity statistics. Contigs assembled from contaminant reads (N = 3832) were identified and filtered from our assembly using the blobtools v1.1 pipeline (Laetsch and Blaxter 2017; RRID:SCR_017618). Jellyfish v2.2.10 (Marcais and Kingford 2012; RRID:SCR_005491) was used for kmer counting (kmer length = 21bp) and the resulting output was used in the GenomeScope2.0 web portal (Ranallo-Benavidez et al. 2020) to estimate genome size (Figure S1). Filtered assembly quality and polishing was carried out using Inspector v1.0.2 (Chen et al. 2021; see Table S1 for details). Gene content completeness was assessed via BUSCO v5.4.7 (Seppey et al. 2019; RRID:SCR_015008) using the Arthropoda_odb10 dataset. Repeat content of the final (filtered and polished) assembly was assessed using RepeatMasker v4.1.3 (Tarailo-Graovac and Chen 2009; RRID:SCR_012954; Table S2) before being structurally annotated with the Helixer v0.3.1 algorithm pipeline (Stiehler et al. 2020; Holst et al. 2023) using the pre-made invertebrate training dataset. Functional annotation was done using eggnog-mapper v5.1.12 (Huertas-Cepas et al. 2019; Cantalapiedra et al. 2021;RRID:SCR_002456).

### Gene family analysis

We selected nine receptor genes associated with nociception when expressed in class III or class IV multidendritic sensory neurons in Arthropoda. This included five sensory receptor channels with GO terms for the detection of thermal, mechanical, and/or chemical stimuli involved in the sensory perception of pain (GO0050965, thermal: pain, TRPA1; GO0050966, mechanical: rpk, pain, ppk, ppk26; GO0050968, chemical: ppk, ppk26, TRPA1), TRPm, Pkd2, and NOMPC (ion channels with a role in cold nociception; Turner et al. 2016), and two mechanoelectrical transduction channels linked to noxious touch sensitization, NOMPC and Piezo (Kim et al. 2012; Hehlert et al. 2021). These seven ortholog families were downloaded from the eggnog database (V5.0, http://eggnog5.embl.de).

Additional species of interest not present in the database (*Hermetia illucens*, BioProject: <u>PRJEB37575</u>; *Tenebrio molitor*, BioProject: <u>PRJNA820846</u>; *Manduca sexta*, BioProject: <u>PRJNA658700</u>; *Penaeus vannamei*, BioProject: <u>PRJNA438564</u>; *Acheta domesticus,* BioProject: <u>PRJNA706033</u>; Bombyx mandarina, BioProject: <u>PRJDB13954</u>; Generalovic et al. 2021, Kaur et al. 2023, Gershman et al. 2021, Zhang et al. 2019, Dossey et al. 2022, Xiang et al. 2018) were manually added to our list by functionally annotating available protein sequences using eggnog-mapper, which uses the eggnog5.0 database to identify orthologous gene families. *Aceta domesticus* did not have publicly available protein sequences, thus we structurally annotated it using Helixer v0.3.1 (Stiehler et al. 2020; Holst et al. 2023) before functional annotation. Phylogenetic analysis (sequence alignment and tree-building) of PAINLESS peptide sequences (Figure 4) was done using Clustal Omega (https://www.ebi.ac.uk/Tools/msa/clustalo/; Madeira et al. 2022; RRID:SCR_001591) before visualization using the ETE TreeView web portal (http://etetoolkit.org/treeview/; Huerta-Cepas et al. 2016). Gene families of interest were also obtained from the newest (v6; Table S3) eggnog database (eggnog6.embl.de; Hernández-Plaza et al. 2023) but additional species were not added.

## Results & Discussion

### *Tenodera sinensis* genome assembly & annotation

The initial assembly was highly fragmented by long-read genome assembly standards (Number of contigs = 8798; N50 = 1.6Mb; Total size = 3.3Gb) and roughly 20% larger than our GenomeScope prediction (2.7Gb; Figure S1). Blobtools analysis determined that the assembly was heavily contaminated with viral and bacterial contigs (N = 3832, Figure S2; 18.81% of raw reads, Table S1). Our contaminant-filtered assembly was still found to be fragmented and larger than expected (Number of contigs = 4966; N50 = 1.8Gb; Total length = 3.03Gb; Table S1). RepeatMasker found that 37.99% of the genome was composed of repetitive elements (Table S2), predominantly retroelements (18.41%) and DNA transposons (18.08%). Inspector analysis found that our assembly had a number of structural errors (N = 593) and small scale errors (N = 83917; 27.7 per Mb), but polishing was able to remove most small-scale errors (after polishing N = 8788; 2.9 per Mb; see Table S1 for details). Despite assembly errors, BUSCO analysis determined that our assembly was highly complete (odb10_insecta Complete = 98.9% [Single copy = 93.5%, duplicated = 5.4%], Fragmented = 0.5%, Missing = 0.6%; Figure 3). Helixer structurally annotated 24,673 genes in our assembly, 14,092 of which were functionally annotated via comparison to the eggnog-mapper database using only orthology to arthropoda. BUSCO assessment of our annotation showed that a substantial portion of total gene content is likely not represented (odb10_insecta Complete = 78.2% [Single copy = 75.1%, duplicated = 3.1%], Fragmented = 7.9%, Missing = 13.9%; Figure 3); however, this was also the case (albeit less drastically) for our helixer annotation of *A. domesticus* (odb10_insecta Complete = 86.7% [Single copy = 82.7%, duplicated = 4.0%], Fragmented = 7.4%, Missing = 5.9%; BUSCO complete in assembly = 94.8%; Dossey et al. 2023) suggesting there may be biases in Helixer’s pre-made invertebrate training dataset which is dominated by insects in the orders diptera and lepidoptera (details regarding helixer training datasets can be found at: https://uni-duesseldorf.sciebo.de/s/lQTB7HYISW71Wi0). Thus, the lower than expected recovery of single copy orthologs may represent this bias, rather than the quality of our assembly.

**Figure 3.**
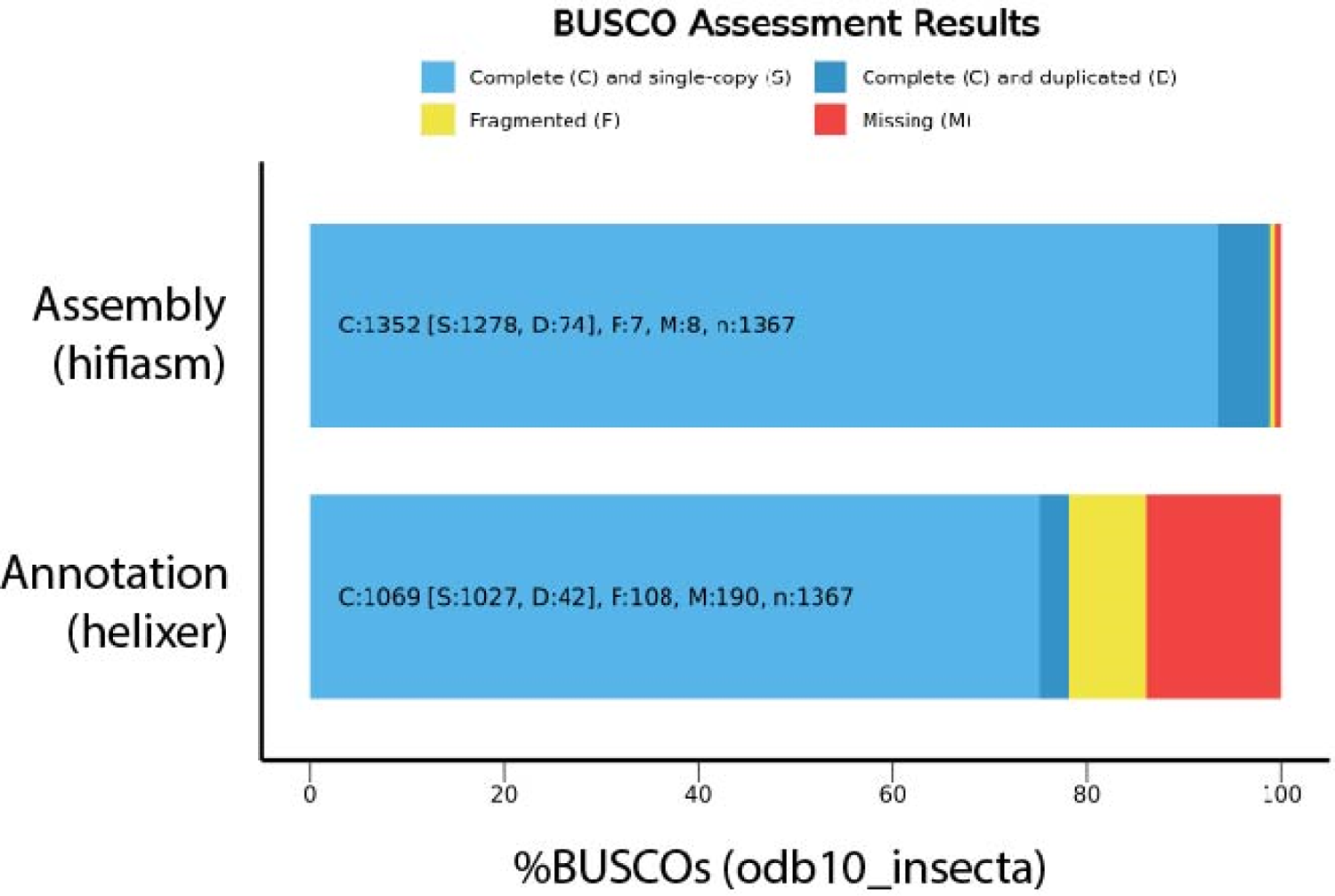
BUSCO assessment of *T. sinensis* assembly and annotation. Annotation assessment was performed in proteome mode.

**Figure 4.**
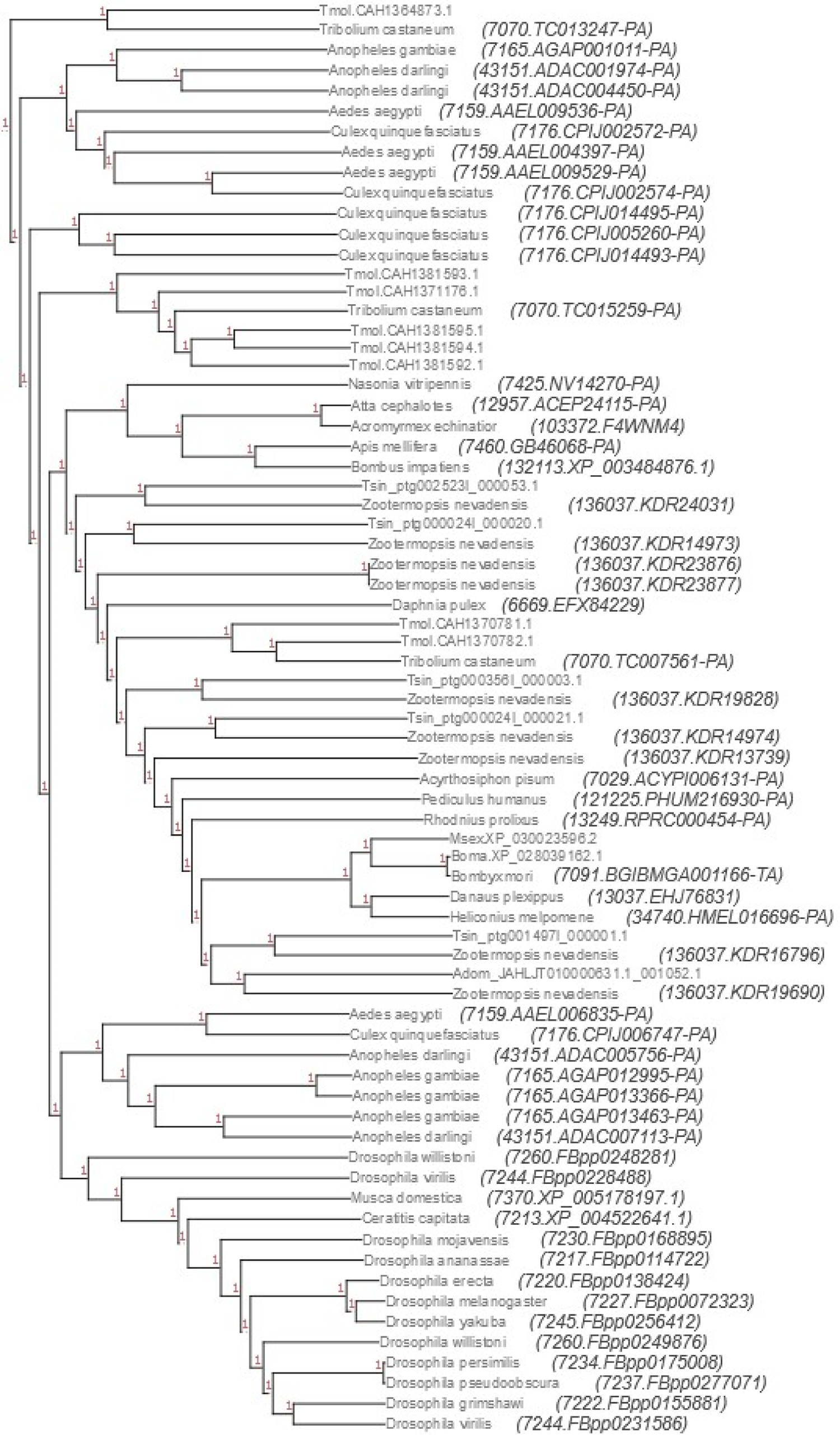
Gene tree showing the relatedness of *painless* genes across our sample of arthropods. *Tenodera sinensis* genes cluster most closely with those of *Zootermopsis nevadensis* (Blattodea). Genes with resolved species names were present in the eggnog5 database (Orthofamily 41TMZ), whereas those with only gene IDs at branch tips are those of our additional species (Adom = *Acheta domesticus*, Boma = *Bombyx mandarina*, Msex = *Manduca sexta*, Tmol = *Tenebrio molitor*, Tsin = *Tenodera sinensis*).

### Variation in ion channel copy numbers in *T. sinensis*

*T. sinensis* was found to have at least one copy of 8 of 9 genes for receptor channels of interest, with the exception being *pkd2* (Table 2). Pkd2 acts as a direct cold sensor in class III md neurons in *D. melanogaster,* and is necessary (alongside NOMPC and TRPm) for aversive responses to cold behavior (Turner et al. 2016). *T. sinensis* was found to have an expansion in both *nompC* and *trpm* copy number (from one each in *D. melanogaster* to three each in *T. sinensis*). The lack of *pkd2* could suggest an inability to directly sense aversive levels of cold in *T. sinensis* despite the presence of *nompC* and *trpm*, but given its function has been characterized solely in *D. melanogaster* and we could not detect an ortholog in many insect genomes, it is difficult to infer this with confidence. Further study of *T. sinensis* behavior and physiology would be needed to assess function. Other orders of insects have evolved novel thermal nociceptors following the loss of other thermosensitive nociceptive ion channel (e.g., duplication and neofunctionalization of Waterwitch into HsTRPA to complement the loss of TRPA1 in the Hymenoptera; Kohno et al. 2010). Further, there is often more than one receptor for assessing potentially noxious thermal information (e.g., the heat sensitivity of Pyrexia, painless, and TRPA1; Tracey et al. 2003, Lee et al. 2007, Wang et al. 2009).

**Table 1.** Ion channel gene families obtained from the EggNog5 database. This table includes gene counts of species included in the database and those obtained from eggnog-mapper outputs of additional species.

|  |  |  |  |  |  |  |  |  |  |
| --- | --- | --- | --- | --- | --- | --- | --- | --- | --- |
|  |  | Orthogroup | 41TMZ | 41U7<br>9 | 41UGV | 41UWY | 41VXR | 41X6F | 41XIN |

| Order | Family | Representative gene name | *pain* | *Nomp C* | *trpm* | *Piezo* | *Pkd2* | *ppk/rpk/ppk26* | *TrpA1* |
| --- | --- | --- | --- | --- | --- | --- | --- | --- | --- |
| Trombidiformes | Tetranychidae | *Tetranychus urticae* | 0 | 2 | 1 | 2 | 1 | 0 | 0 |
| Chilopoda | Geophilomorphae | *Strigamia maritima* | 0 | 1 | 2 | 2 | 1 | 0 | 2 |
| Anomopoda | Daphniidae | *Daphnia pulex* | 1 | 2 | 2 | 0 | 0 | 0 | 0 |
| Decapoda | Penaeidae | *Peneus vannamei* | 0 | 1 | 2 | 2 | 0 | 0 | 2 |
| Blattodea | Archotermopsidae | *Zootermopsis nevadensis* | 9 | 0 | 1 | 1 | 6 | 0 | 1 |
| Mantodea | Mantidae | *Tenodera sinensis* | 5 | 3 | 3 | 2 | 0 | 5 | 1 |
| Psocodea | Pediculidae | *Pediculus humanus* | 1 | 1 | 1 | 3 | 1 | 0 | 1 |
| Hemiptera | Aphididae | *Acyrtosiphon pisum* | 1 | 1 | 2 | 3 | 1 | 3 | 1 |
| Hemiptera | Reduviidae | *Rhodnius prolixus* | 1 | 1 | 2 | 1 | 1 | 1 | 1 |
| Orthoptera | Gryllidae | *Acheta domesticus* | 1 | 4 | 1 | 2 | 0 | 12 | 1 |
| Hymenoptera | Formicidae | *Acromyrmex echinator* | 1 | 4 | 1 | 1 | 0 | 0 | 0 |
| Hymenoptera | Formicidae | *Atta cephalotes* | 1 | 5 | 2 | 1 | 0 | 0 | 0 |
| Hymenoptera | Braconidae | *Microplitis demolitor* | 0 | 2 | 1 | 2 | 0 | 0 | 0 |
| Hymenoptera | Pteromalidae | *Nasonia vitripennis* | 1 | 1 | 1 | 2 | 0 | 0 | 0 |
| Hymenoptera | Apidae | *Apis mellifera* | 1 | 4 | 1 | 1 | 0 | 0 | 0 |
| Hymenoptera | Apidae | *Bombus impatiens* | 1 | 2 | 1 | 1 | 0 | 0 | 0 |
| Coleoptera | Tenebrionidae | *Tribolium castaneum* | 3 | 1 | 1 | 1 | 1 | 17 | 1 |
| Coleoptera | Tenebrionidae | *Tenebrio molitor* | 8 | 1 | 1 | 1 | 1 | 19 | 1 |
| Lepidoptera | Nymphalidae | *Danaus plexippus* | 1 | 1 | 1 | 1 | 0 | 3 | 1 |
| Lepidoptera | Nymphalidae | *Heliconius melpomene* | 1 | 1 | 2 | 1 | 0 | 1 | 1 |
| Lepidoptera | Sphingidae | *Manduca sexta* | 1 | 2 | 8 | 7 | 0 | 4 | 2 |
| Lepidoptera | Bombycidae | *Bombyx mandarina* | 1 | 2 | 5 | 8 | 0 | 3 | 1 |
| Lepidoptera | Bombycidae | *Bombyx mori* | 1 | 1 | 2 | 2 | 0 | 2 | 1 |
| Diptera | Culcidae | *Aedes aegypti* | 4 | 2 | 2 | 1 | 0 | 20 | 2 |
| Diptera | Culcidae | *Anopheles darlingi* | 4 | 1 | 1 | 1 | 0 | 4 | 2 |
| Diptera | Culcidae | *Anopheles gambiae* | 4 | 1 | 1 | 1 | 0 | 6 | 1 |
| Diptera | Culcidae | *Culex quinquefasciatus* | 6 | 2 | 2 | 3 | 0 | 31 | 2 |
| Diptera | Tephritidae | *Ceratitis capitata* | 1 | 1 | 1 | 1 | 2 | 3 | 1 |
| Diptera | Drosophilidae | *Drosophila ananassae* | 1 | 2 | 2 | 1 | 1 | 3 | 1 |
| Diptera | Drosophilidae | *Drosophila erecta* | 1 | 1 | 2 | 1 | 1 | 3 | 1 |
| Diptera | Drosophilidae | *Drosophila grimshawi* | 1 | 1 | 2 | 2 | 3 | 3 | 1 |
| Diptera | Drosophilidae | *Drosophila melanogaster* | 1 | 1 | 1 | 5 | 1 | 3 | 1 |
| Diptera | Drosophilidae | *Drosophila mojavensis* | 1 | 1 | 2 | 2 | 4 | 4 | 1 |
| Diptera | Drosophilidae | *Drosophila persimilis* | 1 | 1 | 2 | 1 | 5 | 4 | 2 |
| Diptera | Drosophilidae | *Drosophila pseudoobscura* | 1 | 1 | 2 | 2 | 5 | 3 | 2 |
| Diptera | Drosophilidae | *Drosophila virilis* | 2 | 1 | 2 | 2 | 3 | 3 | 1 |
| Diptera | Drosophilidae | *Drosophila willistoni* | 2 | 1 | 1 | 2 | 1 | 3 | 1 |
| Diptera | Drosophilidae | *Drosophila yakuba* | 1 | 1 | 2 | 1 | 1 | 3 | 1 |
| Diptera | Muscidae | *Musca domestica* | 1 | 1 | 1 | 2 | 1 | 11 | 2 |
| Diptera | Hermetiinae | *Hermetia illucens* | 0 | 2 | 2 | 1 | 0 | 2 | 6 |

Prior arguments by Eisemann et al. (1984) and others have hypothesized that behavioral non-responsiveness to injury in insects, such as an *T. sinensis* sexual cannibalism, suggests insects are unlikely to perceive (minimally, mechanically) noxious stimuli and thus unlikely to have an adaptive role for pain and sentience. However, since these publications, the genetic components of nociception have been found to be ancient and highly conserved (e.g., Peng et al. 2015); it is, therefore, unsurprising that our data suggest that chemical, mechanical, and thermal nociception are plausible in *T. sinensis.* Still, further tests that confirm 1) these channels are expressed in multidendritic sensory neurons and 2) they have a nociceptive function in the Mantodea will be important for conclusively demonstrating nociception in mantids.

Differential expression and splicing into novel isoforms can lead to different functional roles of these receptors in different cell types (e.g., *dTRPA1;* Zhong et al. 2012, Gu et al. 2022). Amino acid substitution can also lead to variation in channel sensitivity to noxious stimuli, even for orthologous channels (e.g., loss of chemical sensitivity in *Si*HsTRPA versus *Am*HsTRP; Wang et al. 2018). Altogether, these various mechanisms for generating functional plasticity of these receptors in different tissue types suggests moderate caution should be employed when suggesting the presence or absence of a specific gene corresponds to the presence or absence of a specific nociceptive ability in Mantodea. Still, we may broadly expect some perception of noxious stimuli to be highly plausible given this genetic information and the ancient and highly conserved nature of many of these genes. If male mantises are, indeed, able to perceive a mechanically noxious stimulus (such as female cannibalism), this suggests mechanisms other than a lack of nociceptive perception underpin any instances of apparent behavioral unresponsiveness (see Gibbons and Sarlak, 2020). Instead, our genetic data supports prior behavioral evidence that shows male mantises attempt to avoid being cannibalized (Lelito and Brown, 2006; Brown et al., 2012) and provides further support for the ancient and highly conserved nature of the peripheral components of pain pathways across Arthropoda.

### Variation in ion channel copy number across Arthropoda

All the gene families except the *ppk/rpk/ppk26* family saw reasonably similar ranges of copy number across the surveyed arthropods (from 0 up to 6 - 9 copies). However, greater variation (0 - 31 copies) was found in the *ppk/rpk/ppk26* family, with roles in mechanical and chemical nociception. This included substantial intra-order variation (e.g., 2 copies in *Hermetia illucens,* 11 copies in *Musca domestica,* and 31 copies in *Culex quinquefasciatus*, all dipterans). Prior research on mosquitoes has found that additional ion channel gene copies play a role in non-nociceptive sensory processes, specifically heat-perception involved with prey location (Wang et al. 2009). Our results suggest that ion channel copy numbers are highly variable and that they are likely to play an important role in the evolution of Arthropod sensory systems, but more precise studies of expression dynamics and cellular functions are necessary to better understand this process.

Our search also specifically focused on insects that are mass-reared (such as the black soldier fly, *H. illucens*) or even domesticated (such as the silkworm moth, *Bombyx mori*) to see if there were any patterns of ion channel loss or duplication associated with the novel ecological conditions posed by rearing at scale on farms under significant human management. Generally, farmed species retained at least one copy of all genes except *pkd2,* with only *Tenebrio molitor* and *M. domestica* having a copy of this gene. This could suggest, as in *T. sinensis,* some loss of cold nociception capabilities that should be confirmed functionally (and considered alongside the aforementioned caveats).

The only other gene loss event was *painless* in *H. illucens;* however, *H. illucens* had significant duplication of *trpa1* (6 copies), where no other arthropod species in our study had more than two copies of this gene. It is possible the additional copies of *trpa1* may serve to recover some of the lost functions of *painless* as both are TRPA proteins that have roles, for instance, in thermal nociception in other Diptera (Hwang et al. 2012; Neely et al. 2011). *T. molitor* had significant duplication of *painless* (8 copies) and *ppk/rpk/ppk26* (19 copies). *Acheta domesticus* also had significant duplication of *ppk/rpk/ppk26* (12 copies). *B. mori* had significant duplication of *piezo* (8 copies) and *trpm* (5 copies). These genes were both highly expressed in the two lepidopterans - *B. mori* and *Manduca sexta* - but not in any other order we reviewed. Overall, these results suggest that domestication and/or mass rearing do not result in nociceptive ion channel loss in insects, which means it will still be important to monitor and reduce the occurrence of noxious stimuli in mass rearing facilities as part of ethical mini-livestock husbandry and slaughter.

### Concluding Remarks

In summary, we have produced a high-quality draft genome of the Chinese mantis (*Tenodera sinensis*) and used it to identify multiple putative nociceptive ion channels.

Comparative analyses suggest that some of these genes have undergone multiple duplications that are shared with Blattodea, whereas others have been lost in these closely related clades. These findings suggest that mantids are likely capable of perceiving different types of noxious stimuli (extreme heat/cold, mechanical damage, etc.). Future research into their precise functional roles will elucidate the evolution of nociception in an iconic sexually cannibalistic species.

## Acknowledgments

Thanks to Alexander Glica and Andrew Fairclough for helping to rear mantises, and Chris Wirth for taking our voucher specimens.

## Supplementary Figures

**Figure S1.**
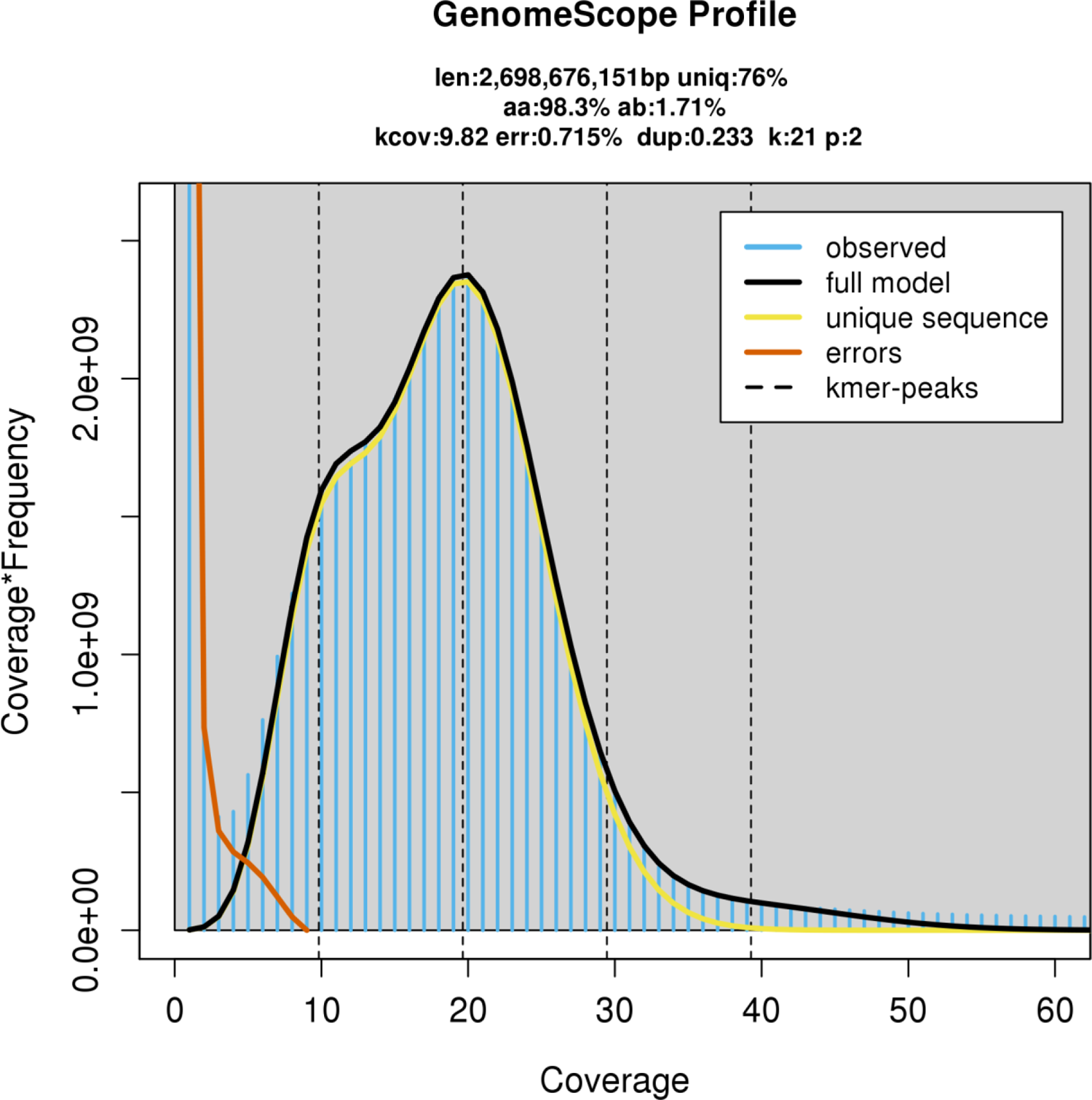
Genomescope profile plot showing that our assembly is heterozygous and estimated to be 2.7Gb in size, similar to other sequenced mantid genomes.

**Figure S2.**
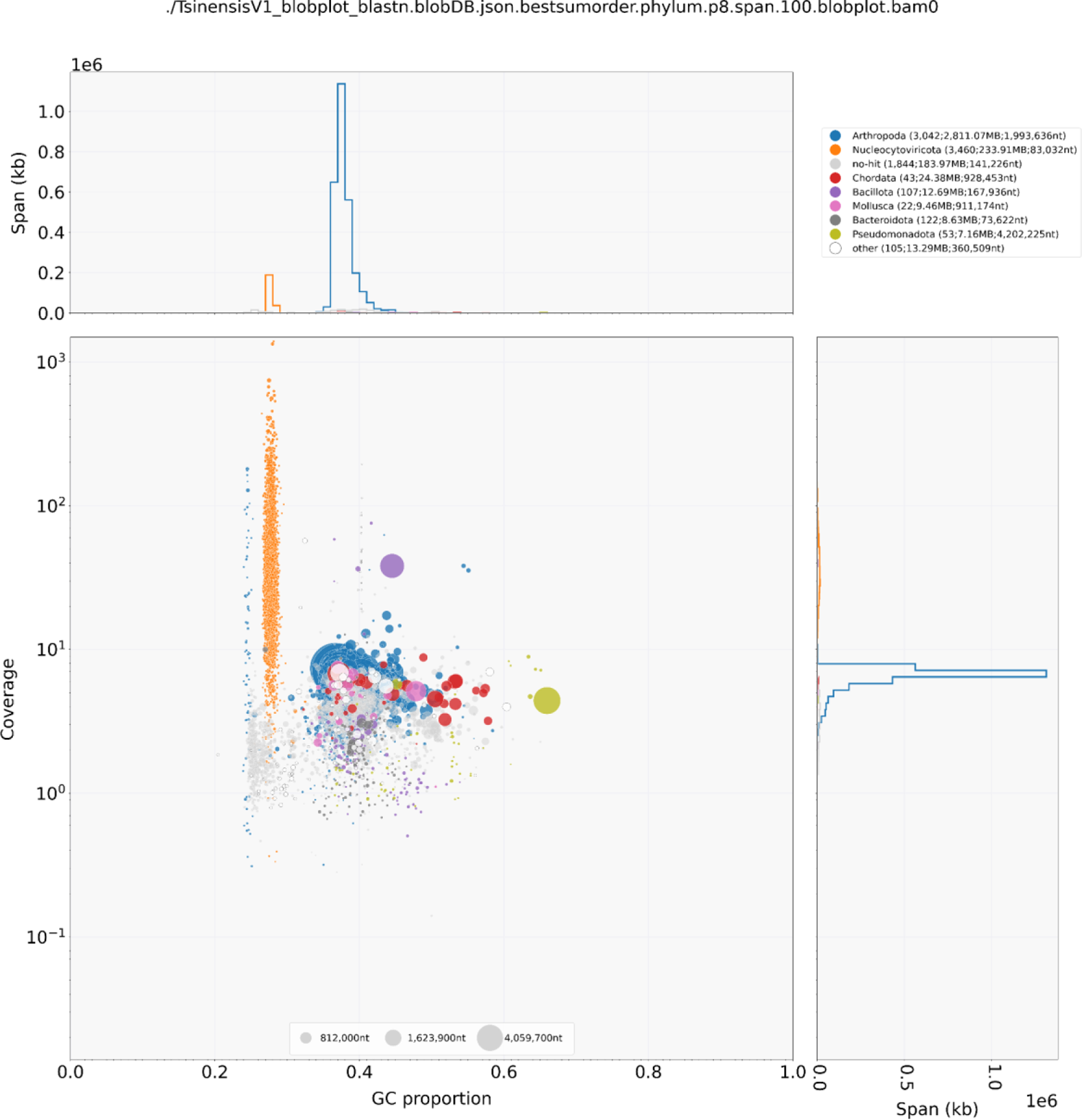
Blobplot showing the presence of substantial viral and bacterial contamination in our genome assembly. Contigs identified as viral or bacterial (N = 3832) were filtered out of our assembly.

---

**Table S1.** Summary statistics of genome assembly before and after polishing with Inspector.

|  | Before | After |
| --- | --- | --- |
| Number of contigs | 4966 | 4966 |
| Number of contigs > 10000 bp | 4966 | 4966 |
| Number of contigs >1000000 bp | 837 | 837 |
| Total length | 3032411541 | 3032411605 |
| Total length of contigs > 10000 bp | 3032411541 | 3032411605 |
| Total length of contigs >1000000bp | 2084261305 | 2084260901 |
| Longest contig | 16238473 | 16238500 |
| Second longest contig length | 14940332 | 14940332 |
| N50 | 1806180 | 1806165 |
| N50 of contigs >1Mbp | 1806180 | 1806165 |
| Mapping rate /% | 81.19 | 81.19 |
| Split-read rate /% | 7.83 | 7.83 |
| Depth | 25.541 | 25.5411 |
| Mapping rate in large contigs /% | 61.28 | 61.28 |
| Split-read rate in large contigs /% | 6.64 | 6.63 |
| Depth in large conigs | 28.1542 | 28.1532 |
| Structural error | 593 | 606 |
| Expansion | 386 | 391 |
| Collapse | 172 | 177 |
| Haplotype switch | 31 | 34 |
| Inversion | 4 | 4 |
| Small-scale assembly error /per Mbp | 27.67335464 | 2.898023469 |
| Total small-scale assembly error | 83917 | 8788 |
| Base substitution | 61937 | 7186 |
| Small-scale expansion | 10715 | 703 |
| Small-scale collapse | 11265 | 899 |
| QV | 38.4297348 | 39.15840409 |

**Table S2.**
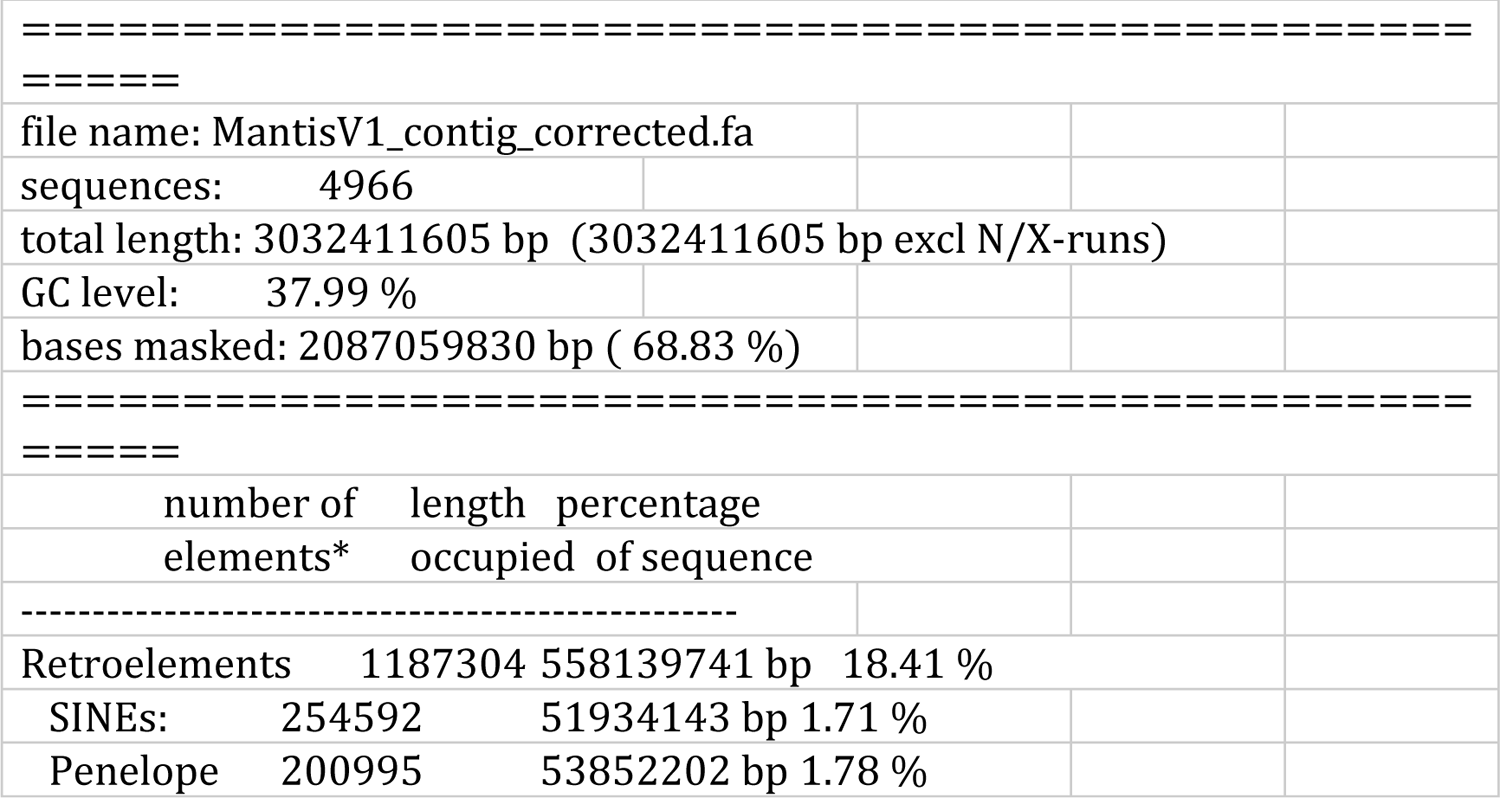
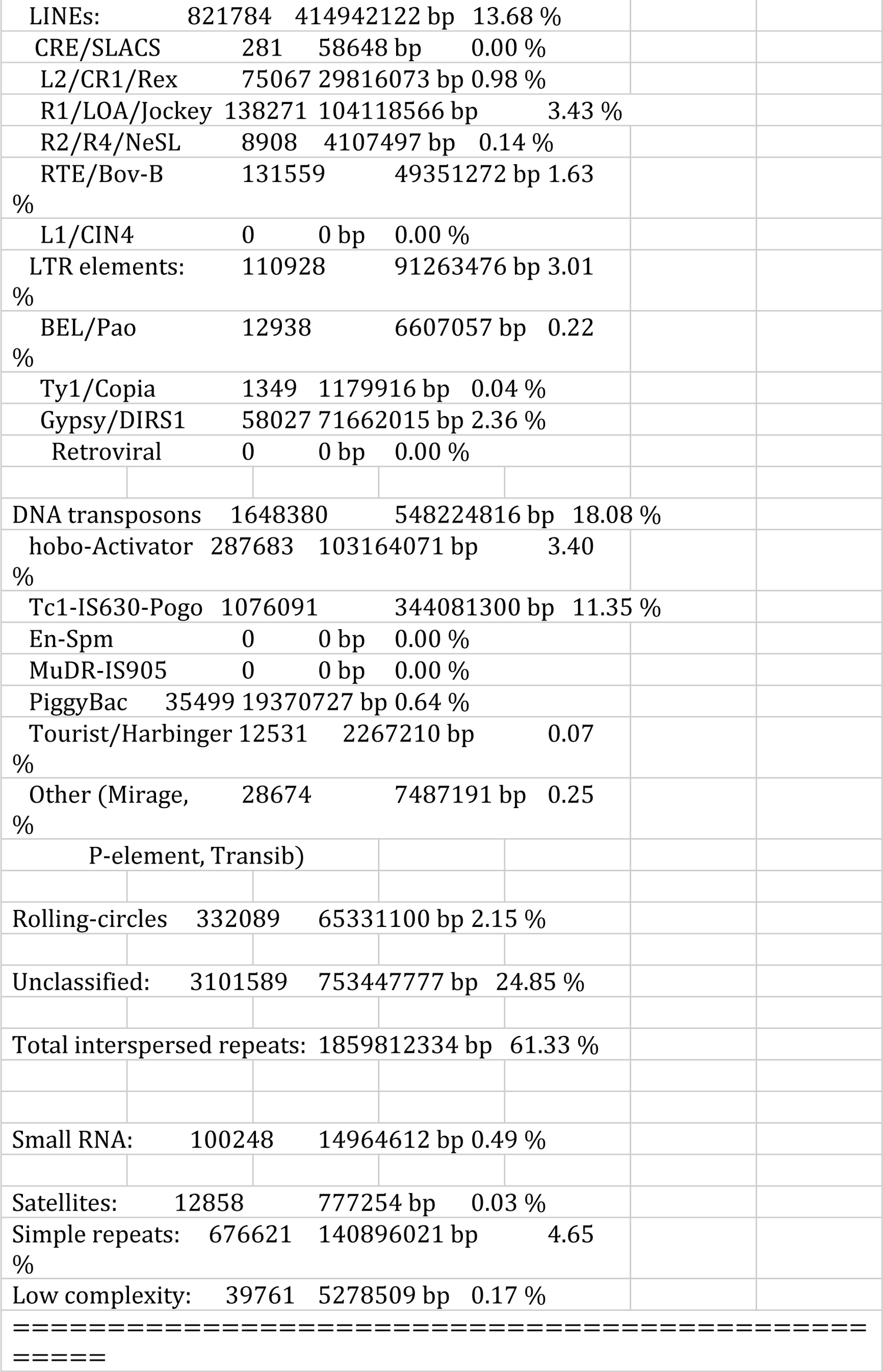
Summary of GC content and repetitive element content from RepeatMasker.

**Table S3.**
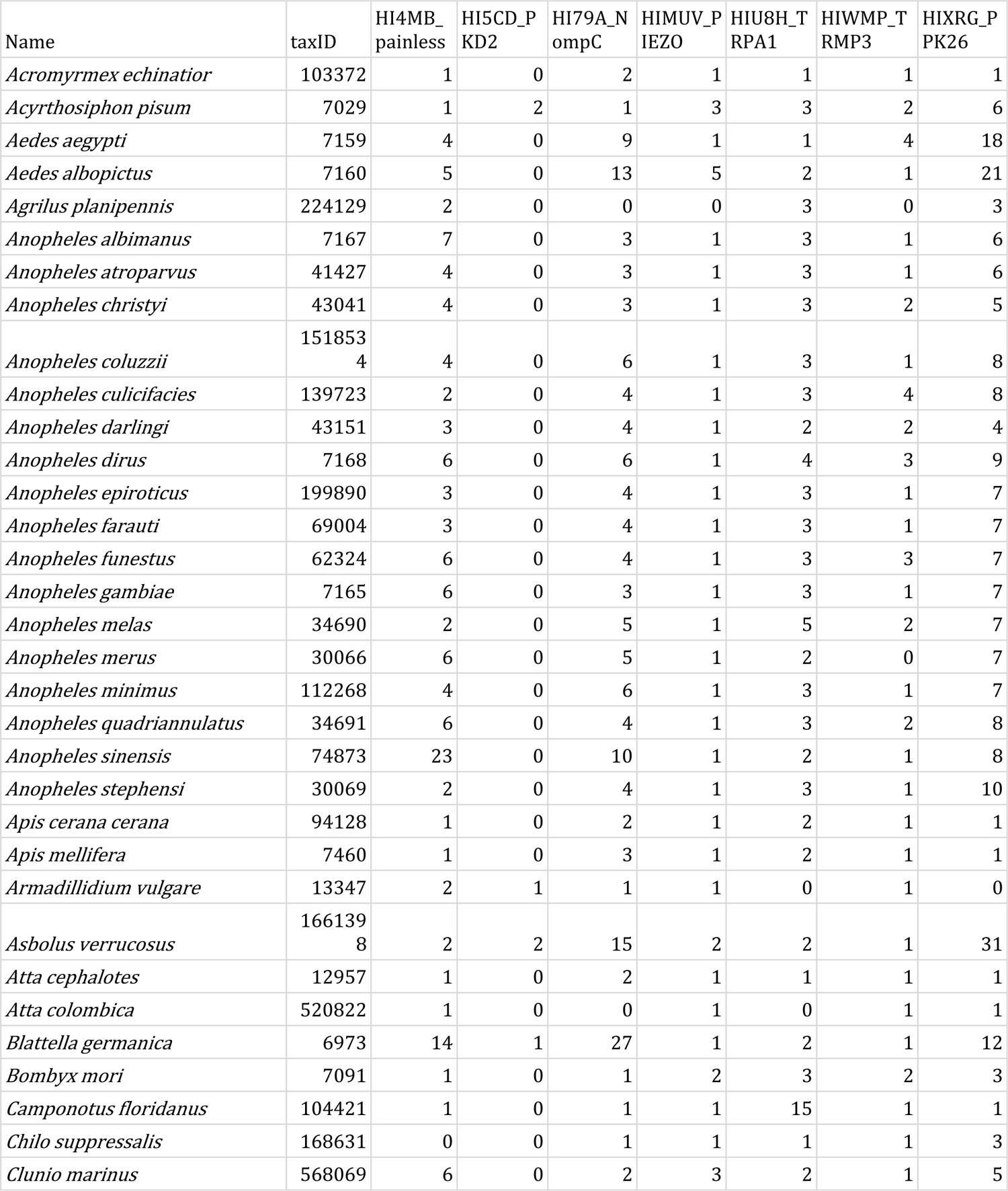
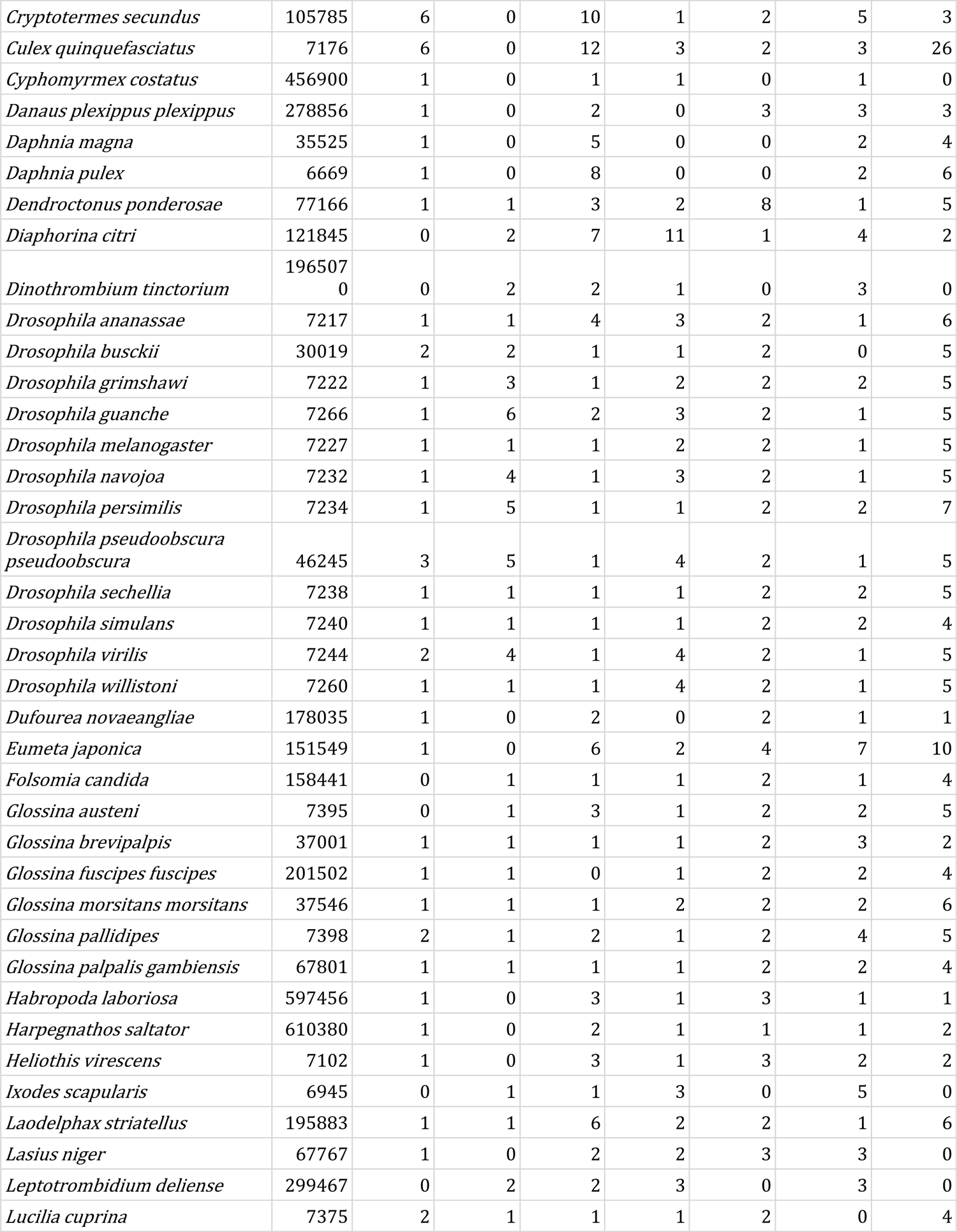
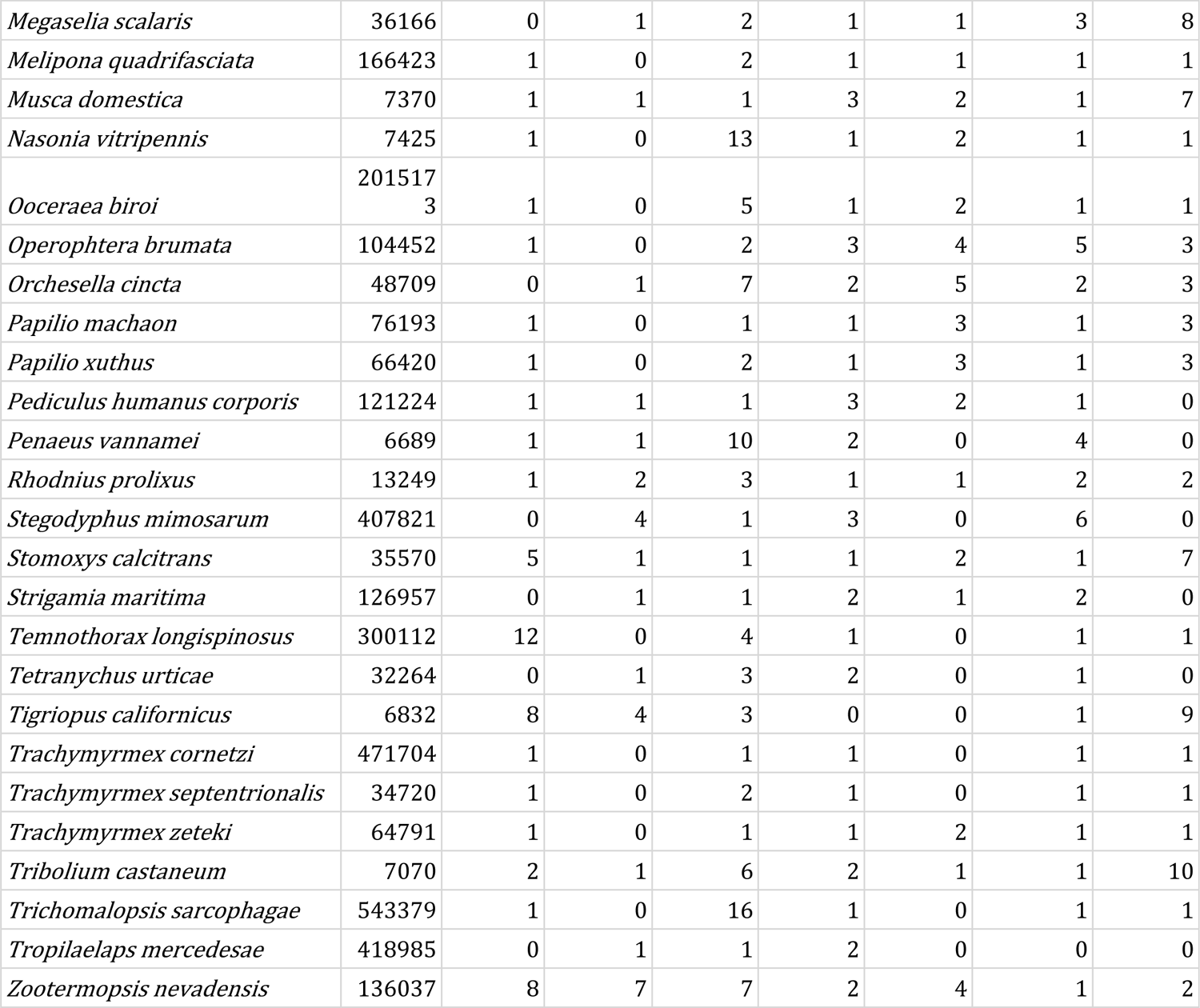
Ion channel gene families obtained from the EggNog6 database. Only genes present within the database are included.

